# KCNQ channels enable reliable presynaptic spiking and synaptic transmission at high-frequency

**DOI:** 10.1101/2019.12.26.888875

**Authors:** Yihui Zhang, Dainan Li, Youad Darwish, Laurence O. Trussell, Hai Huang

**Author notes:** Address correspondence to: Hai Huang, Ph.D., Department of Cell and Molecular Biology, Tulane University, 2000 Percival Stern Hall, 6400 Freret St, New Orleans, LA 70118 USA.

## Abstract

The presynaptic action potential (AP) results in calcium influx which triggers neurotransmitter release. For this reason, the AP waveform is crucial in determining the timing and strength of synaptic transmission. The calyx of Held nerve terminals of rat show minimum changes in AP waveform during high-frequency AP firing. We found that the stability of the calyceal AP waveform requires KCNQ K^+^ channel activated during high-frequency spiking activity. High-frequency presynaptic spikes gradually led to accumulation of KCNQ channels in open states which kept interspike membrane potential sufficiently negative to maintain Na^+^ channel availability. Accordingly, blocking KCNQ channels during stimulus trains led to inactivation of presynaptic Na^+^, and to a lesser extent K_V_1 channels, thereby reducing the AP height and broadening AP duration. Thus, while KCNQ channels are generally thought to prevent hyperactivity of neurons, we find that in axon terminals these channels function to facilitate high-frequency firing needed for sensory coding.

**HIGHLIGHTS:**

- KCNQ channels are activated during high-frequency firing
- The activity of KCNQ channels helps the recovery of Na^+^ and K_V_1 channels from inactivation and maintains action potential waveform
- Reliable presynaptic action potential waveform preserves stable Ca^2+^ influx and reliable synaptic signaling

## INTRODUCTION

At chemical synapses, the presynaptic action potential (AP) drives the activation of voltage-gated Ca^2+^ channels, and its repolarization enhances the driving for Ca^2+^ influx into the terminal. Together these two factors determine the intracellular Ca^2+^ levels needed for triggering vesicle fusion and neurotransmitter release (Borst and Sakmann, 1996; Sabatini and Regehr, 1999). Thus, the size and shape of the presynaptic AP is a key determinant of timing and strength of synaptic transmission (Sabatini and Regehr, 1997; Geiger and Jonas, 2000; Hoppa et al., 2014). Due to the differential expression of diverse voltage-dependent ion channels, the waveforms and firing patterns of action potentials vary considerably among various types of neurons in the mammalian brain (Bean, 2007). Moreover, the AP waveform is subject to short- and long-term regulation. For example, serotonin inhibits presynaptic potassium channels and broadens the action potential in Aplysia, which leads to increased calcium influx and enhanced neurotransmitter release (Siegelbaum et al., 1982). During repetitive firing of action potentials, voltage-gated Na^+^ (Na_V_) and K^+^ (K_V_) channels undergo cycles of activation and deactivation, inactivation and repriming. Slow cumulative inactivation of voltage-gated ion channels under repetitive high-frequency firing may affect the size and waveform of AP. For example, the AP waveforms of the pituitary nerve terminals and the mossy fiber boutons are altered and the duration of action potentials is substantially increased during moderate to high-frequency firing (Jackson et al., 1991; Geiger and Jonas, 2000).

The globular bushy cells of the cochlear nucleus faithfully relay auditory signals to the medial nucleus of the trapezoid body (MNTB) principal neurons through a giant glutamatergic synapse, the calyx of Held (Mc Laughlin et al., 2008; Lorteije et al., 2009). Reliability and precision of synaptic transmission are required in the calyx-MNTB synapse in order to encode location of transient sounds in natural environments (Joris and Trussell, 2018). One factor that contributes to this precision is the remarkable stability of the presynaptic waveform during ongoing firing (Sierksma and Borst, 2017). Indeed, calyx of Held terminals show only minor AP waveform change at spike frequencies up to hundreds of Hertz. However, the biophysical basis of this constancy is unclear. We explored the role of presynaptic KCNQ (K_V_7) channels. These channels are thought to temper spike activity in neurons, as naturally occurring mutations in KCNQ genes are associated with seizure, and pharmacological blockade of KCNQ channels leads to neuronal hyperactivity (Biervert et al., 1998; Peters et al., 2005; Brown and Passmore, 2009; Qi et al., 2014). Here, we found that KCNQ channels in calyx nerve terminals are cumulatively activated during high-frequency AP activity, thus minimizing inactivation of Na^+^ and K_V_1 channels, and maintaining a stable presynaptic AP waveform. Blocking KCNQ channels disrupts the Ca^2+^ influx, reduced the synaptic transmission, and disrupted the high-fidelity synaptic signaling. These results illustrate how a slowly gating K^+^ channel is essential to enable high-frequency firing needed for sensory coding.

## RESULTS

### KCNQ channels regulate the terminal intrinsic excitability

Using recordings from the calyx nerve terminal, we confirmed previous observations made in neuronal somata that block of KCNQ channels results in hyperactivity, when assayed by delivering long depolarizing current pulses. Typically, calyces responded to long current pulses by generating only a few action potentials at the current onset, regardless of the length of the pulse (Fig. 1) (Dodson et al., 2003; Ishikawa et al., 2003). Loss of KCNQ conductance had a striking effect on presynaptic excitability. Application of XE991 (10 μM), a concentration that specifically blocks KCNQ channel in the MNTB (Huang and Trussell, 2011), caused calyces to fire continuously and at high rates for the duration for the stimulus over 1 s current injections of 50-200pA (n = 6; Fig. 1B, G). The KCNQ channel activator flupirtine (10 μM) however decreased spike number during current injections (n = 4; Fig. 1E, H). Due to partial activation of KCNQ channels at rest (Huang and Trussell, 2011), application of either drug caused a shift in resting potential (see below). Therefore, after applying the drugs and recording responses to current pulses, we then injected positive or negative bias current to restore the membrane potential back to the control resting level and repeated the current pulse protocol. Similar results were obtained (Fig. 1C, F-H). Together, these show that excitability of the terminal is dependent not only on the fast-gating channels that underlie the action potential, but also on slow conductances like KCNQ.

**Figure 1.**
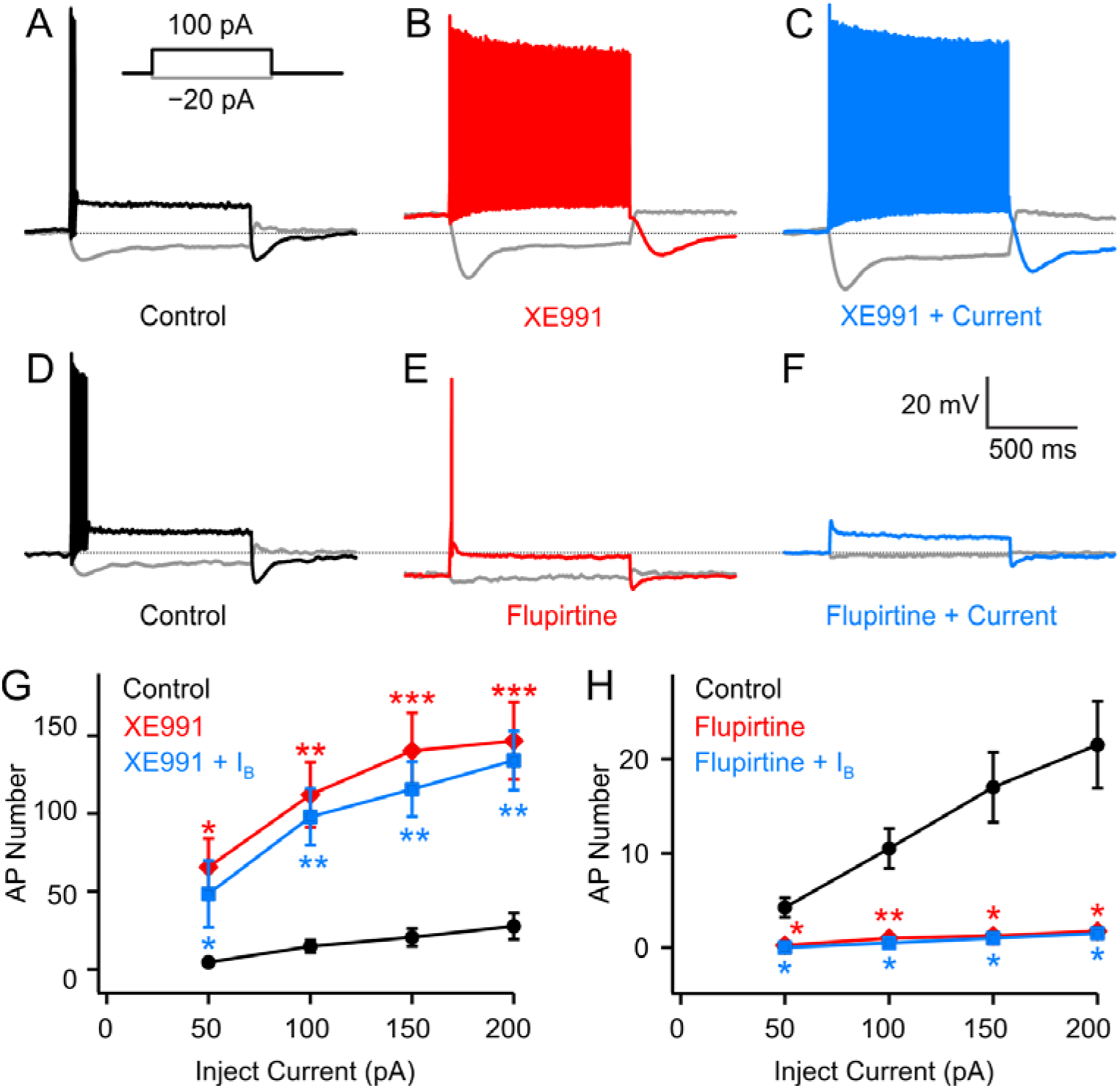
KCNQ channels suppress firing during prolonged current injection. (A-B) APs elicited by 1-s current steps in control conditions (A) and after (B) bath application of 10 μM XE991. (C) The excitability change was not restored when bias currents were injected to correct for the depolarization of RMP. (D-F) Similar to A-C, except bath application of 10 μM flupirtine to open KCNQ channels. (G-H) AP numbers elicited by 1-s depolarizing current steps to the values given on the x-axis before and after bath application of XE991 (G) and flupirtine (H). XE991 increased, while flupirtine decreased, the total number of APs elicited by the 1 s-long depolarizing current steps. *P < 0.05, **P < 0.001, ***P < 0.001; paired Student’s t-test. Error bars are mean ± S.E.M.

### KCNQ channels maintain AP waveform during high-frequency firing

We next asked how blocking KCNQ channels affected conducted, rather than locally triggered action potentials. To do that we first stimulated presynaptic axons and assessed the stability of the AP waveform when evoked at different stimulus rates. The AP waveforms were remarkably stable during prolonged repetitive stimulation (400 spikes) at high frequencies (Fig. 2). AP waveforms remained nearly unchanged during 10 and 33 Hz firing. Increasing the firing frequency to 100 Hz induced a slight amplitude decrease and AP broadening in activity-dependent. Comparing the 400^th^ AP to the 1^st^ AP, the amplitude decreased by 3.0 ± 0.9% and half-width increased by 12.9 ± 2.9%. When the stimulation frequency increased to 333 Hz, AP amplitude decreased by 21.2 ± 3.6% and the half-width increased by 76.4 ± 20.1% (n = 5). Thus, the calyceal terminals have the capacity to maintain stable AP waveform during high-frequency firing.

**Figure 2.**
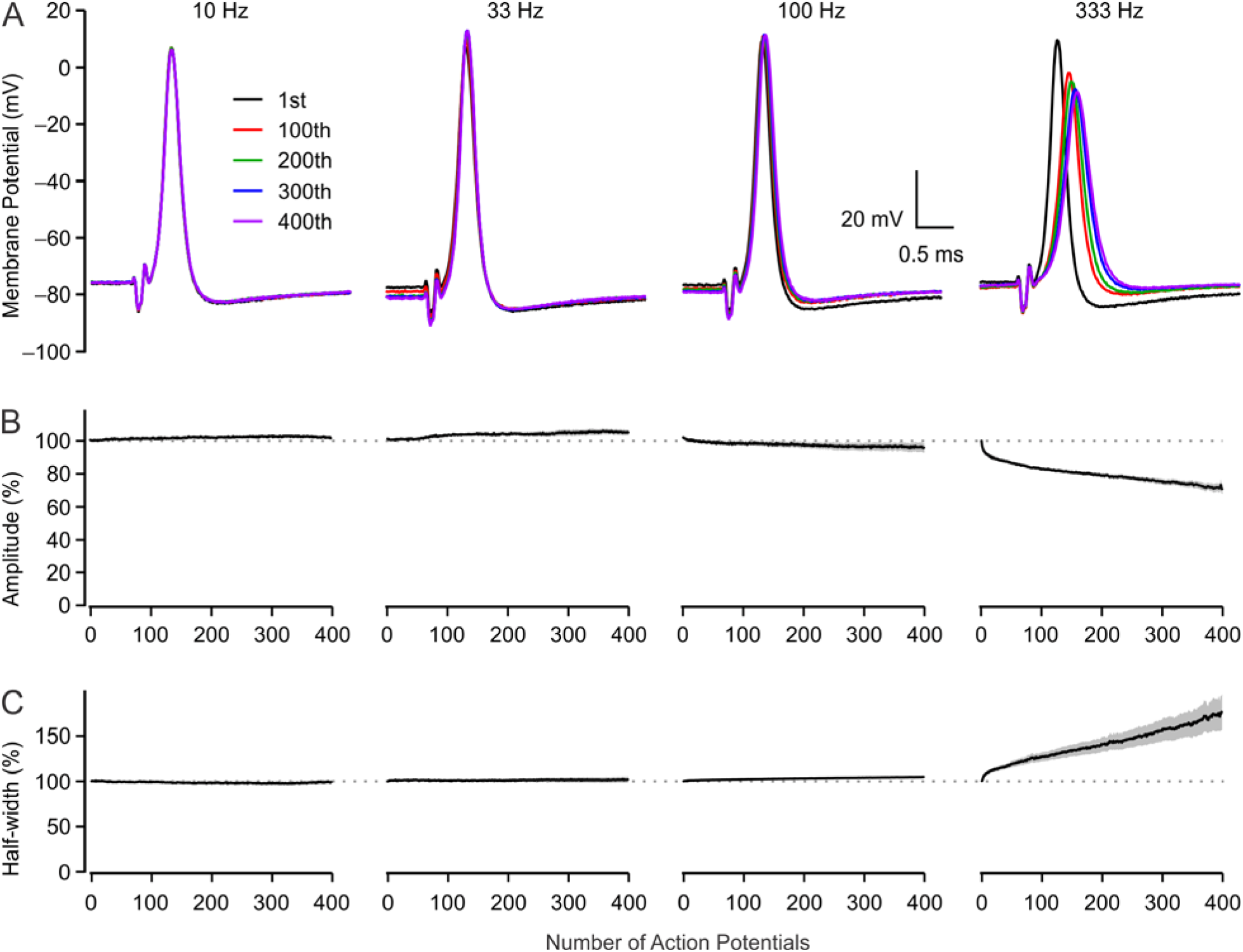
Stable AP waveform during high-frequency AP firing in calyx of Held. (A) Trains of APs evoked at the frequency of 10, 33, 100 and 333 Hz by afferent fiber stimulation. Every 100^th^ AP is superimposed with the first AP. (B-C) Plots of AP amplitude (B) and half-width (C) during spiking at each frequency. Data were normalized to the value of the first AP in each train.

We then investigated the underlying mechanisms that support reliable firing at the calyx terminal using 333-Hz stimulation protocol. When measuring the inter-spike potential (ISP), i.e., the afterhyperpolarization membrane potential between spikes (0-0.2 ms before the next stimulation artifact), we noticed that 6 out of 10 calyces exhibited a depolarization during repetitive firing during the first 20-50 ms of the stimulus protocol, and then gradually hyperpolarized (Fig. 3A,F, black trace). The overall hyperpolarization of ISP was 3.2 ± 0.7 mV (n = 6). We hypothesized that a slow K^+^ current, whose kinetics is similar to KCNQ channels (Huang and Trussell, 2011), is activated during high-frequency spiking that is critical in maintaining the reliable AP waveform. Indeed, in the presence of 10 μM XE991, the delayed hyperpolarization of ISP was eliminated. Instead, the ISP continuously depolarized during high-frequency spiking (Fig. 3B, F, red trace). Accompanying the depolarization of ISP, the AP waveforms were greatly altered. When the first and last APs in the spike train were compared, it was apparent that XE991 had a significantly greater impact on the waveform of the last rather the first APs. The amplitude of the last AP was reduced by 46.6 ± 5.5% (P = 0.002, comparing to the control), and the half-width was broadened to 299 ± 45% (P = 0.02; n = 5) after application of XE991 (Fig. 3). KCNQ channels in the calyx of Held have an activation threshold more hyperpolarized than the resting membrane potential (RMP) and thus contribute to set the resting properties of the calyx of Held (Huang and Trussell, 2011). Blocking KCNQ channels with XE991 depolarized the membrane potential by about 5 mV, an effect that impacts the activation and inactivation of other voltage-gated channels. After applying XE991, we injected a hyperpolarizing bias current to restore the membrane potential, and repeated the stimulating protocol. The waveform of the first APs were restored to the control level, while the late phase was not fully restored: the ISP continued depolarizing during the whole stimulation (Fig. 3C,F) and the AP became shorter and broader. Overall, the AP height was reduced to 57.6 ± 4.9% (P = 0.01) and half-width was broadened by 251 ± 12% (Fig. 3; P = 0.003; n = 5). Another KCNQ blocker, linopirdine, had a similar effect (n = 4, data not shown). These results indicate the accumulated activation of KCNQ channels during high-frequency AP firing produce a gradual hyperpolarization of ISP, which is essential in maintaining AP waveform during high-frequency firing.

**Figure 3.**
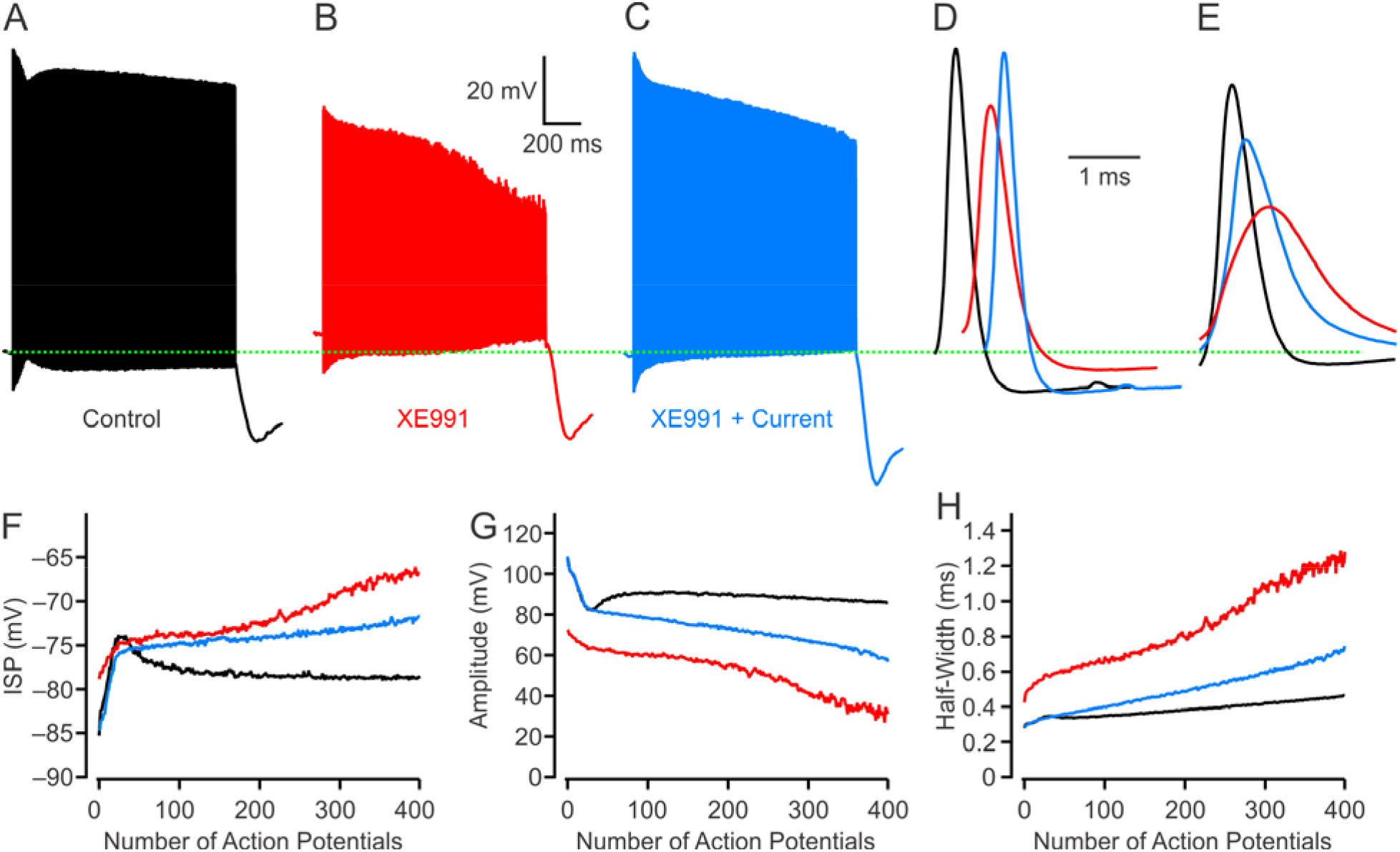
KCNQ channels contribute to reliable AP waveform during high-frequency firing. (A-C) Trains of APs elicited at the frequency of 333 Hz (3-ms intervals) before (black, A) and after (red, B) bath application of 10 μM XE991. Bias current was injected to restore the RMP to the value of control AP train (blue, C). (D-E) The first (D) and 400^th^ (E) AP were superimposed at an expanded time scale. (F-H) Plots of inter-spike potential (ISP, F), height (G), and half-width (H) of AP waveforms shown in A-C.

### Activation of KCNQ prevents accumulative inactivation of Na^+^ and K_V_1 channels

During repetitive firing, Na_V_ and K_V_ channels undergo cycles of activation and deactivation, inactivation and repriming (recovery from inactivation). Repriming is strongly dependent on the spike afterpotential (Raman and Bean, 2001; Bean, 2007). Since activation of KCNQ channels keeps the ISP hyperpolarized, we hypothesized that the hyperpolarization induced by opening of KCNQ channels during high-frequency firing helps Na_V_ and K_V_ channels to recover from inactivation. To test whether KCNQ channels influences the recovery of Na^+^ from inactivation during high-frequency (400 APs at 333 Hz) firing, we recorded presynaptic Na_V_ and K_V_ currents before and after AP trains under voltage-clamp in the presence of blockers for other voltage-gated channels (see Experimental Procedures). The spikes recorded in control (Fig. 3A), with XE991 (Fig. 3B), and XE991 with tonic bias current (Fig 3C) were used as voltage command templates. 10-ms depolarizing square pulses from resting membrane potential to −10 mV were applied before and immediately after the AP command train (3-ms after the onset of the last AP) (Fig. 4A). The ratio of the peak inward current evoked by the second over the first square pulse represents the degree of inactivation developed during the train stimulation. The Na^+^ current was reduced to 65.8% ± 1.2% after the control AP train template. When templates were used from spikes recorded in the presence of XE991 or XE991 plus bias current conditions, the inactivation was greatly increased. The peak inward Na^+^ current decreased to 26.6% ± 2.1% (P < 0.001) and 25.8% ± 2.1% (P < 0.001), respectively (Fig. 4B-D; n = 8), indicating that KCNQ channels serve to minimize the cumulative inactivation of Na^+^ channels during repetitive firing.

**Figure 4.**
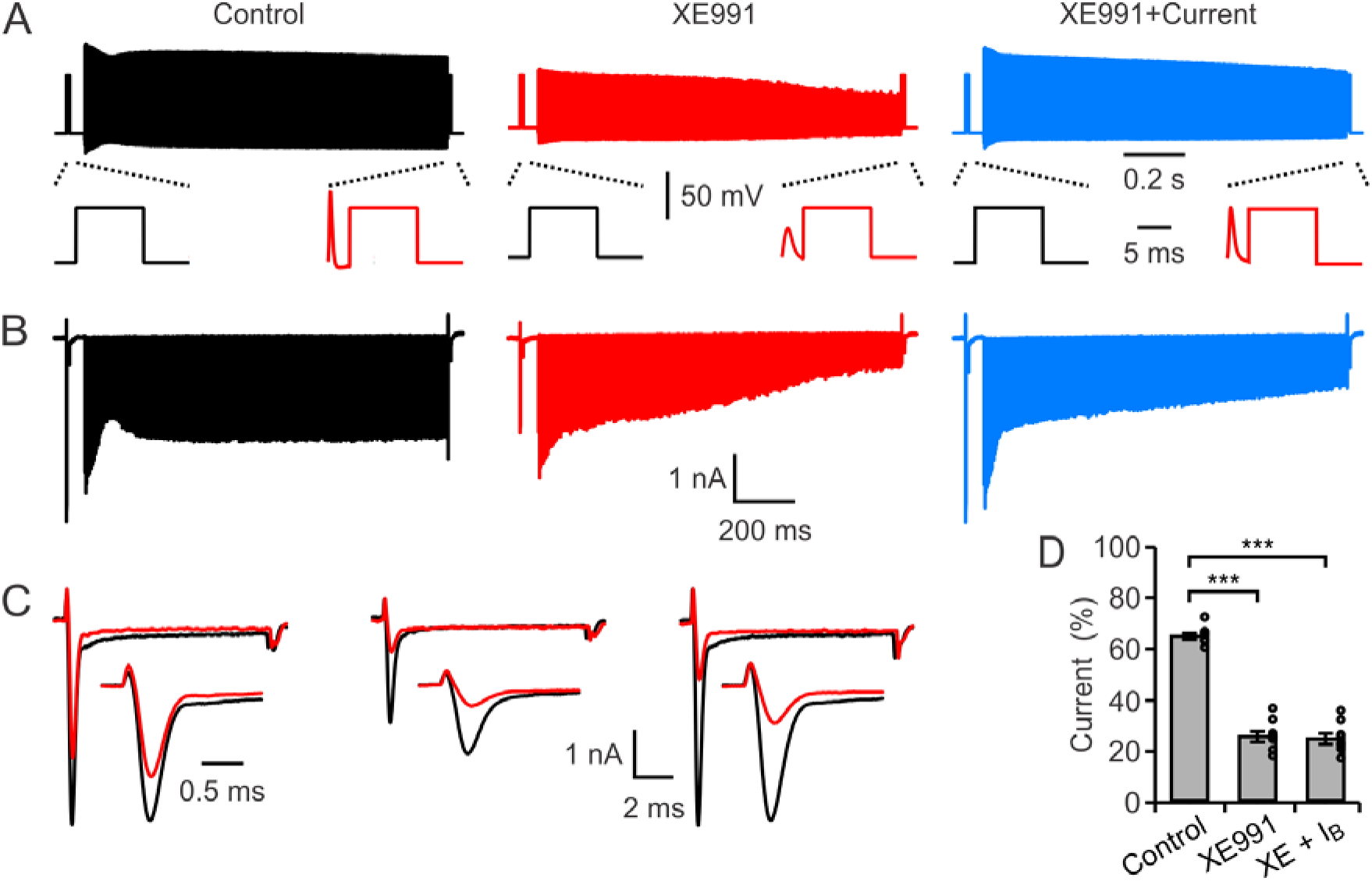
KCNQ activation relieves accumulative inactivation of Na^+^ channels at high-frequency firing. (A) Command templates were created by adding 10-ms depolarizing pulses before and immediately after AP trains obtained under control, XE991, and XE991 + current conditions shown in Fig 3A-C. (B) Na^+^ current (I_Na_) evoked by the command templates. (C) I_Na_ evoked by the depolarizing pulses (10 ms depolarization to −10 mV) before (black) and after (red) AP train were superimposed at an expanded time scale. (D) Summarizing plot of the Na^+^ current ratio, showing that blocking XE991 enhanced Na^+^ channel inactivation. Recordings were made in the presence of TEA, 4-AP, XE991, Cd^2+^ and Cs^+^ to block the voltage-gated K^+^, Ca^2+^, and HCN, respectively.***P < 0.001, Student’s t-test. Error bars are mean ± SEM.

Similar experiments were used to test the inactivation of K_V_1 and K_V_3 channels, which are the predominant K^+^ channel subtypes in the calyx of Held (Dodson et al., 2002; Ishikawa et al., 2003; Dodson and Forsythe, 2004; Song et al., 2005). K_V_1 currents (measured in the presence of 1 mM TEA) were reduced by repetitive spike activity to 64.1% ± 3.6% under control spike train, and further reduced to 60.5% ± 2.6% and 54.9% ± 2.7% under XE991 and XE991 plus bias current conditions, respectively (Fig. 5A-D), indicating a slightly larger inactivation after blocking KCNQ channels (n = 5; P = 0.04 for control vs. XE991; P < 0.01 for control vs. XE991 plus bias current). K_V_3 current were recorded in the presence of K_V_1 channels blocker margatoxin (10 nM). In control conditions, K_V_3 currents were reduced to 50.1% ± 6.1%, likely due to the use-dependent inactivation (Marom and Levitan, 1994), which was not difference from XE991 conditions (51.4% ± 6.1%) or XE991 + current conditions (50.2% ± 6.1%) (Figure 5E-G; P = 0.66 and 0.96, respectively; n = 7). Therefore, the gradual hyperpolarization of the membrane during repetitive firing primarily serves to minimize inactivation of Na^+^ channels.

**Figure 5.**
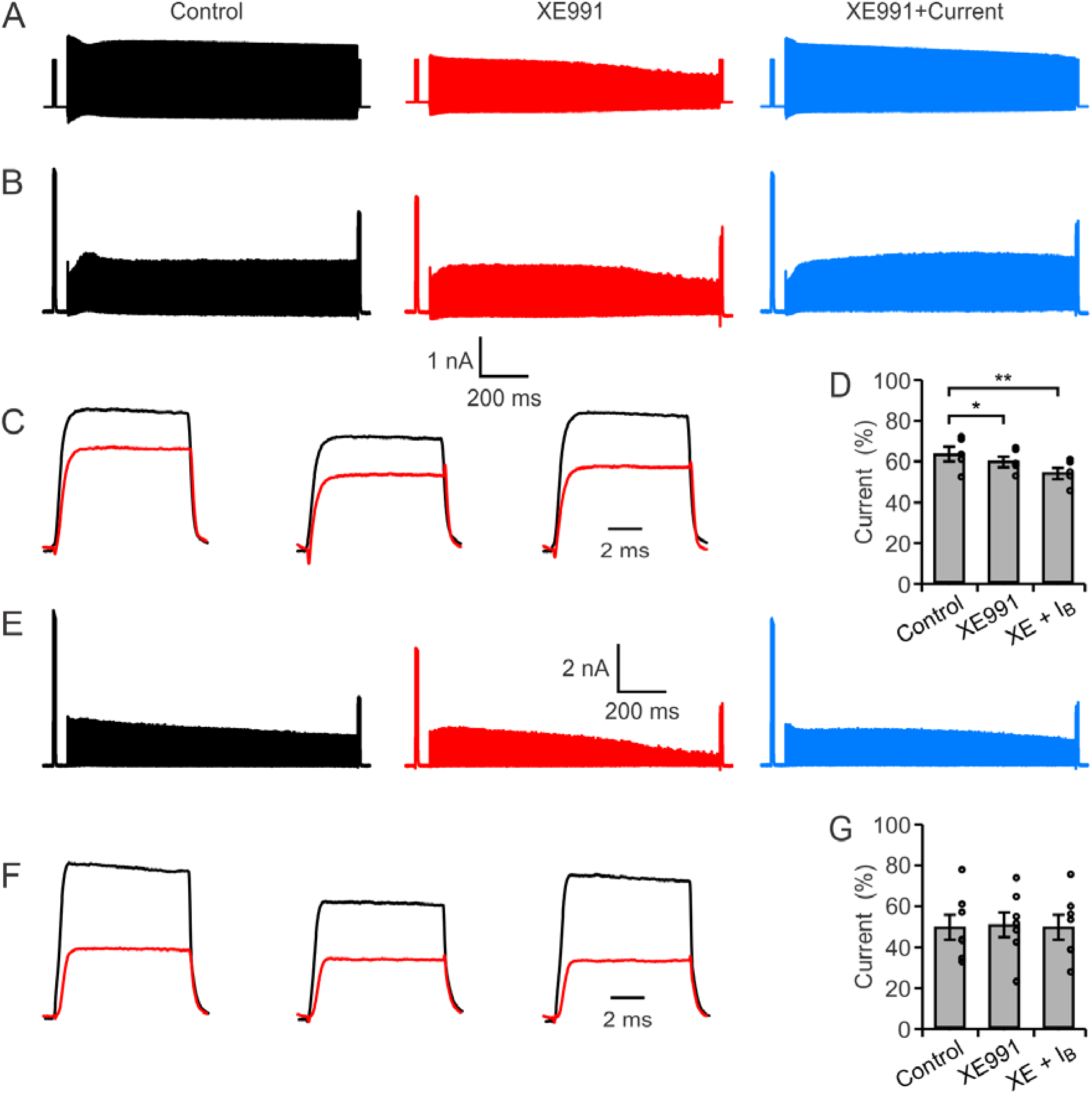
KCNQ activation relieves accumulative inactivation of Kv1, but not Kv3. (A) Recordings were similar to Fig. 4 while K^+^ current were recorded. (B) Representative traces of K_v_1 current evoked by the command templates. Recordings were made in the presence of TTX, Cd^2+^, Cs^+^, XE991 and TEA to block Na^+^, Ca^2+^, HCN, KCNQ, and Kv1, respectively. (C) I_Kv1_ evoked by the 1^st^(Black) and 2^nd^ (Red) depolarizing pulses (10 ms depolarization to −10 mV) were superimposed at an expanded time scale. (D) Summarizing data of the remaining I_Kv1_ for the control, XE991, and XE991 + current command templates. (E-G) Similar to B-D, except K_V_3 currents were recorded in the TTX, Cd^2+^, Cs^+^, XE991 and margatoxin to block Na^+^, Ca^2+^, HCN, KCNQ, and K_V_3, respectively. **P < 0.01; *P < 0.05, Student t-test. Error bars are mean ± SEM.

### KCNQ channels enable reliable calcium influx during high-frequency firing

Presynaptic AP directs Ca^2+^ influx required for vesicle fusion and neurotransmitter release (Sabatini and Regehr, 1997; Geiger and Jonas, 2000; Hoppa et al., 2014). We then tested how changes in AP waveform during high-frequency firing affects Ca^2+^ current. AP trains recorded during current-clamp (Fig. 2) were used as command templates to evoke Ca^2+^ current (I_Ca_) under voltage-clamp recordings (Fig. 6A-B). In control conditions, the I_Ca_ amplitudes were relatively stable, reducing slightly from 2.00 ± 0.14 nA at the first AP to 1.75 ± 0.07 nA at 400^th^ APs. However, I_Ca_ amplitudes were reduced even more, from 1.2 ± 0.09 nA to 0.43 ± 0.06 nA in XE991 conditions; and from 1.87 ± 0.10 nA to 1.43 ± 0.15 nA for the conditions of XE991 + current (Fig 6E). Since the I_Ca_ duration was altered (Fig. 6C-D), we also calculated the AP-induced Ca^2+^ charge. The charge was slightly increased from 0.51 ± 0.05 pC to 0.67 ± 0.06 pC in control conditions, mainly due to AP broadening. In XE991, the AP Ca^2+^ charge was initially 0.45 ± 0.05 pC, briefly increased during the train, and then declined to 0.37 ± 0.06 pC. In XE991 + bias current, Ca^2+^ charge was gradually increased from 0.50 ± 0.06 pC to 0.86 ± 0.08 pC (P < 0.0001). Together, these results indicate that KCNQ channels during high-frequency firing have profound effects on regulating the I_Ca_ and the KCNQ-enabled stable AP waveform is crucial for stably evoking Ca^2+^ influx.

**Figure 6.**
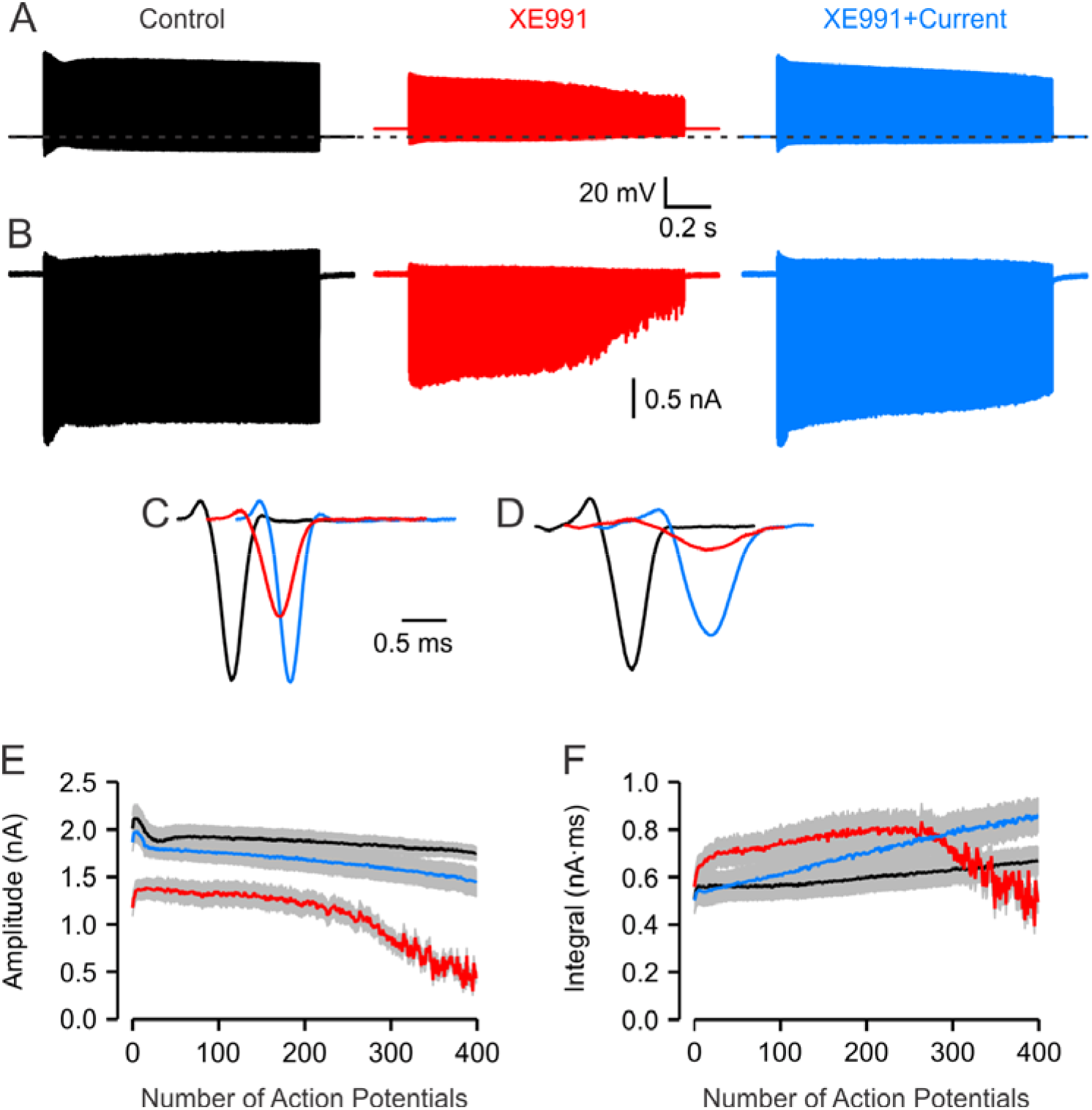
KCNQ activation maintains reliable Ca^2+^ influx during high-frequency firing. (A) Naive control, XE991, and XE991 + current AP trains were used as voltage-clamp command templates (B) Representative traces of I_Ca_ evoked by three command templates with the same calyx. (C-D) First (C) and last (D) I_Ca_ in each recording are superimposed at an expanded time scale. (E-F) amplitudes (F) and integrals (E) of I_ca_ were measured and plotted against the number of the APs within the three command templates.

### KCNQ channels enable reliable synaptic transmission at high-frequency

As blocking KCNQ channels affect the AP-evoked Ca^2+^ influx, we tested the role of KCNQ channels in neurotransmitter release and synaptic transmission. Postsynaptic cells were voltage clamped and presynaptic axons were stimulated with an extracellular electrode to evoke 400 excitatory postsynaptic currents (EPSCs) at 3-ms intervals. Despite synaptic depression, EPSCs were evoked in response to each presynaptic AP (Fig. 7A, B). Application of 10 μM XE991 increased the EPSC amplitudes at the beginning of the train due to the previously described role of KCNQ channels on presynaptic resting membrane potential (Huang and Trussell, 2011). However, as the AP train proceeded, the EPSCs became less reliable, and littered with small and asynchronous neurotransmitter release events (Fig. 7C,D). The amplitude of 351^st^−400^th^ EPSCs was reduced from 294±125 pA to 259±131pA (P = 0.03, n = 5, paired t-test). High-frequency globular bushy cell signals are faithfully transmitted to the postsynaptic MNTB neurons through the calyx terminal and each presynaptic spike triggers an AP at its postsynaptic MNTB neuron, with few failures (Mc Laughlin et al., 2008; Lorteije et al., 2009). We therefore tested whether KCNQ contributes to faithful one-to-one synaptic signaling. Postsynaptic cells were current clamped to record the AP firing in response to the presynaptic stimulation. At 333 Hz, each presynaptic stimulation evoked an AP in MNTB neuron under control condition (Fig. 7E,G). Bath application with XE991 did not affect the reliability of synaptic transmission at the beginning of the train, but eventually disrupted one-to-one fidelity and caused failures in postsynaptic spikes (Fig. 7F, G). Overall, the failure rate increased from 5.7 ± 4.2% to 31.7 ± 6.5% in the 351-400^th^ stimulations (P = 0.0009, n = 7), indicating that the activation of presynaptic KCNQ current during high-frequency firing is critical in controlling reliable neurotransmitter release and synaptic signaling across the synapse.

**Figure 7.**
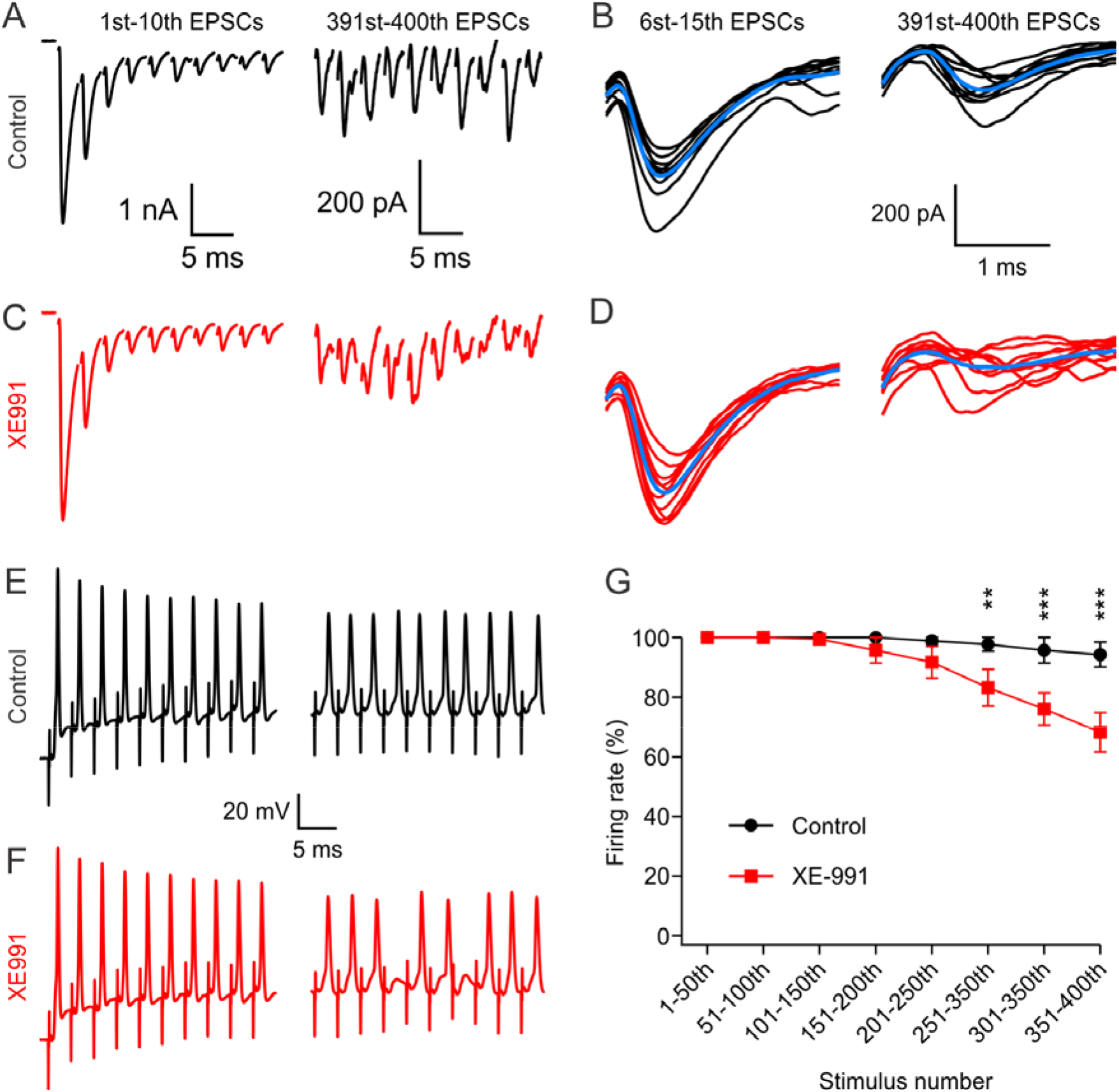
Blocking of KCNQ channels reduces the reliability of synaptic transmission at high frequency. (A) EPSCs evoked by 1^st^−10^th^ (left) and 391^st^−400^th^ (right) AP at 333 Hz. (B) The 6^th^−15^th^ EPSCs and 391^st^−400^th^ EPSCs were superimposed (average shown as blue). (C,D) The same EPSC recordings in the presence of 10-20 μM KCNQ. (E,F) Representative trace showed the same range of number of APs before (black) and after (red) bath application of XE991. (G) Statistical results showed that XE991 decreased the reliable synaptic transmission at the late phase during high-frequency synaptic transmission.

## DISCUSSION

Calyx of Held terminals exhibit minimum changes in AP waveform during high-frequency spiking. Here, we showed that KCNQ channels, most likely KCNQ5 homomers (Huang and Trussell, 2011), are cumulatively activated during high-frequency AP firing, and this activation keeps the presynaptic membrane potential between spikes hyperpolarized, which minimizes cumulative inactivation of Na^+^ and Kv1 channels. The resulting consistency in the AP waveform ensures reliable calcium influx and glutamate release.

### KCNQ channels and reliable AP waveform

While the somata of globular bushy cell and postsynaptic MNTB neurons may exhibit significant AP depression during brief high-frequency firing (Smith and Rhode, 1987; Lorteije et al., 2009; Zhang and Huang, 2017), the AP waveform of the calyceal terminals was remarkably stable during high-frequency firing. In P12–P14 rat brain slices, reliable synaptic transmission was generally possible at frequencies up to 600 Hz for 50 stimuli (Taschenberger and von Gersdorff, 2000). *In* vivo recordings from mice showed that AP showed little or no (<4%) depression when instantaneous firing frequencies over 200 Hz (Sierksma and Borst, 2017). Here we showed amplitude decrease during 100 Hz firing for 4 s was 3% while 333 Hz firing decreased the AP amplitude by 20% (Fig. 2), indicating very stable AP waveforms during prolonged high-frequency firing. Two factors appeared to be crucially important for the lack of a change in the AP’s shape: fast recovery of Na^+^ channels from AP depression (Leao et al., 2005) and relative stable while negative membrane potential following the AP (Sierksma and Borst, 2017). The recovery of Na^+^ from inactivation is sharply dependent on the membrane potential, as 20 mV of hyperpolarization double the speed of recovery (Leao et al., 2005). Because of its slow activation and deactivation kinetics, the amplitude of KCNQ current gradually increases during repetitive APs. This accumulation of KCNQ current hyperpolarized the afterpotential and speeds the recovery of Na^+^ channel from inactivation. Blocking KCNQ channels elevated the membrane potential between action potentials, and led to a distortion of the presynaptic Ca^2+^ current. Therefore, activation of KCNQ channels during high-frequency activity enable the AP firing with stable waveform and high-fidelity synaptic signaling.

### KCNQ channels and control of firing properties

The KCNQ (Kv7) channels are a subfamily of voltage-gated K^+^ channels that are widely expressed in the central nervous system. The slow activation, and non-inactivation properties of KCNQ channels make it suitable for a variety of roles associated with controlling excitability in the brain. In particular, extensive studies reported that KCNQ channels function to prevent excessive activity in various neurons. KCNQ mutation or pharmacological block of KCNQ channels elevated neuronal excitability (Soh et al., 2014; Martinello et al., 2019). *In vivo* studies showed that injection of XE991 leads to hyperactivity (Schwarz et al., 2006), and mutation in KCNQ channels are associated with epilepsy (Biervert et al., 1998). Consistent with these observations, we found that KCNQ blockers caused calyces to fire continuously and at high rates in response to prolong current injection.

However, such analyses do not address the role of K^+^ channels in maintaining the membrane in a spike-ready state, in particular with a membrane potential appropriate for optimal repriming of Na^+^ channels. Loss of Na^+^ channel availability is particularly acute during high-frequency AP activity, and thus a mechanism is needed to ensure stable responses across different activity rates. KCNQ channels are well-suited to this role. At rest, a small proportion of KCNQ channels are activated (Wladyka and Kunze, 2006; Huang and Trussell, 2011; Hu and Bean, 2018), contributing to establishing the resting potential and the baseline responsiveness of soma, axon, and synapse. However, we show here that during repetitive activity, gradual depolarization of the membrane reduces availability of Na^+^ channels, and to a lesser extent Kv1 channels. This loss of channel function broadens and attenuates the spike. At the nerve terminal these changes are particularly problematic because of their effect on Ca^2+^ flux and Ca^2+^-dependent exocytosis.

Although KCNQ channels activate slowly upon depolarization (Biervert et al., 1998; Peretz et al., 2010; Huang and Trussell, 2011), fast spikes will contribute slightly but incrementally to a gradual enhancement of the KCNQ current, and help maintain the membrane potential between spikes at an optimal level. This effect will be of particular importance in axons and terminals of sensory pathways, where driven spikes rates can reach hundreds of Hertz in response to strong stimuli. Thus, besides their role in preventing hyperexcitability, KCNQ channels may function to enable sensory coding across the full dynamic range of the system.

## METHODS

### Slice Preparation

The handling and care of animals were approved by the Institutional Animal Care and Use Committee of Tulane University and in compliance with U.S. Public Health Service guidelines. Brainstem slices containing the MNTB were prepared from P10-16 Wistar rats of either sex as previously described (Huang and Trussell, 2014; Zhang and Huang, 2017). Briefly, 210 μm sections were cut in ice-cold, low-Ca^2+^, low-Na^+^ saline using a vibratome (VT1200S, Leica), incubated at 32°C for 20–40 min in normal artificial cerebrospinal fluid (aCSF) and thereafter stored at room temperature before experiments. The saline for slicing contained (in mM) 230 sucrose, 10 or 25 glucose, 2.5 KCl, 3 MgCl_2_, 0.1 CaCl_2_, 1.25 NaH_2_PO_4_, 25 NaHCO_3_, 0.4 ascorbic acid, 3 myo-inositol, and 2 Na-pyruvate, bubbled with 5% CO_2_/95% O_2_. The aCSF for incubation and recording contained (in mM) 125 NaCl, 10 or 25 glucose, 2.5 KCl, 2 CaCl_2_, 1 MgCl_2_, 1.25 NaH_2_PO_4_, 25 NaHCO_3_, 0.4 ascorbic acid, 3 myo-inositol, and 2 Na-pyruvate, pH 7.4 bubbled with 5% CO_2_/95% O_2_.

### Whole-Cell Recordings

Brain slices were transferred to a recording chamber and were continually perfused with aCSF (2–3 ml/min) warmed to ~32°C by an inline heater (Warner Instruments). Neurons were viewed using an Olympus BX51 microscope with a 40X water-immersion objective and customized infrared Dodt gradient contrast optics. Whole-cell current- and voltage-clamp recordings were made with a Multiclamp 700B amplifier (Molecular Devices).

For current-clamp recordings, pipette solution contained (in mM) 135 K-gluconate, 10 KCl, 4 MgATP, 0.3 Tris-GTP, 7 Na_2_-phosphocreatine, 0.2 EGTA, 10 HEPES, (290 mOsm, pH 7.3 with KOH). For K^+^ current recordings, pipette solution contained (in mM) 135 K-gluconate, 10 KCl, 4 MgATP, 0.3 Tris-GTP, 7 Na_2_-phosphocreatine, 0.2 EGTA, 10 HEPES, (290 mOsm, pH 7.3 with KOH). TTX (0.5 μM), CdCl_2_ (100 μM), CsCl (2 mM), and XE991 (10 μM) were added to the recording solution to block the Na^+^, Ca^2+^, HCN, and KCNQ channels, respectively. For Na^+^ and Ca^2+^ current recordings, pipette solution included (in mM): 120 Cs-methanesulfonate, 20 TEA-Cl, 1 MgCl_2_, 10 HEPES, 5 EGTA, 0.4 Tris-GTP, 3 Mg-ATP, and 5 Na_2_-phosphocreatine (290 mOsm, pH 7.3 with CsOH). To isolate Na^+^ currents in response to AP-voltage command templates, CdCl_2_ (100 μM), TEA (10 mM), 4-aminopyridine (2 mM), XE991 (10 μM), and CsCl (2 mM) were added to block the Ca^2+^, K^+^, HCN, respectively. For Ca^2+^ current recordings, TTX (0.5 μM), CsCl (2 mM), TEA (10 mM), 4-aminopyridine (2 mM), and XE991 (10 μM) were added to the recording solution to block the Na^+^, HCN, and K^+^ channels, respectively. Equimolar NaCl was reduced to keep the osmolarity. For EPSC recordings, pipette solution contained (in mM) 130 cesium methanesulfonate, 10 CsCl, 10 HEPES, 5 EGTA, 0.3 Tris-GTP, 4 Mg-ATP and 5 Na_2_-phosphocreatine (290 mOsm, pH 7.3 with CsOH). Strychnine (1 μM), picrotoxin (50 μM), and (R)-CPP (5 μM) were added to the recording solution to block glycine, GABA, and NMDA receptor–mediated currents.

Pipettes pulled from thick-walled borosilicate glass capillaries (WPI) had open tip resistances of 2–4 MΩ when filled with the above pipette solutions. Series resistances (4– 15 MΩ) were compensated by 60%–80% (bandwidth 3 kHz). Signals were filtered at 4-20 kHz and sampled at 10-50 kHz. Liquid junction potentials were measured and adjusted appropriately.

### Drugs

Drugs were obtained from Alomone labs (XE991), Abcam (TTX), and all others from Sigma. Drugs were stored as aqueous stock solutions at −20°C and dissolved the stock solutions in aCSF to the final concentration immediately before experiments.

### Analysis

Data were analyzed using Clampfit (Molecular Devices), Igor (WaveMetrics). Statistical significance was established using paired and unpaired t-tests as indicated. Data are expressed as mean ± S.E.M.

## AUTHOR CONTRIBUTIONS

Y.Z., Y.D., D.L., and H.H. performed experiments, Y.Z. and H.H. analyzed experiments, Y.Z., L.O.T. and H.H. designed experiments and wrote the manuscript.

## ACKNOWLEDGEMENTS

We thank Dr. Laura Schrader for critical reading of the manuscript. Financial supported was provided by U.S. National Institutes of Health grant R01DC016324 to H. Huang.

## REFERENCES

Bean BP (2007) The action potential in mammalian central neurons. Nat Rev Neurosci 8:451–465.

Biervert C, Schroeder BC, Kubisch C, Berkovic SF, Propping P, Jentsch TJ, Steinlein OK (1998) A potassium channel mutation in neonatal human epilepsy. Science 279:403–406.

Borst JG, Sakmann B (1996) Calcium influx and transmitter release in a fast CNS synapse. Nature 383:431–434.

Brown DA, Passmore GM (2009) Neural KCNQ (Kv7) channels. Br J Pharmacol 156:1185–1195.

Dodson PD, Forsythe ID (2004) Presynaptic K+ channels: electrifying regulators of synaptic terminal excitability. Trends Neurosci 27:210–217.

Dodson PD, Barker MC, Forsythe ID (2002) Two heteromeric Kv1 potassium channels differentially regulate action potential firing. J Neurosci 22:6953–6961.

Dodson PD, Billups B, Rusznak Z, Szucs G, Barker MC, Forsythe ID (2003) Presynaptic rat Kv1.2 channels suppress synaptic terminal hyperexcitability following action potential invasion. J Physiol 550:27–33.

Geiger JR, Jonas P (2000) Dynamic control of presynaptic Ca(2+) inflow by fast-inactivating K(+) channels in hippocampal mossy fiber boutons. Neuron 28:927–939.

Hoppa MB, Gouzer G, Armbruster M, Ryan TA (2014) Control and plasticity of the presynaptic action potential waveform at small CNS nerve terminals. Neuron 84:778–789.

Hu W, Bean BP (2018) Differential Control of Axonal and Somatic Resting Potential by Voltage-Dependent Conductances in Cortical Layer 5 Pyramidal Neurons. Neuron 97:1315–1326 e1313.

Huang H, Trussell LO (2011) KCNQ5 channels control resting properties and release probability of a synapse. Nat Neurosci 14:840–847.

Huang H, Trussell LO (2014) Presynaptic HCN channels regulate vesicular glutamate transport. Neuron 84:340–346.

Ishikawa T, Nakamura Y, Saitoh N, Li WB, Iwasaki S, Takahashi T (2003) Distinct roles of Kv1 and Kv3 potassium channels at the calyx of Held presynaptic terminal. J Neurosci 23:10445–10453.

Jackson MB, Konnerth A, Augustine GJ (1991) Action potential broadening and frequency-dependent facilitation of calcium signals in pituitary nerve terminals. Proc Natl Acad Sci U S A 88:380–384.

Joris PX, Trussell LO (2018) The Calyx of Held: A Hypothesis on the Need for Reliable Timing in an Intensity-Difference Encoder. Neuron 100:534–549.

Leao RM, Kushmerick C, Pinaud R, Renden R, Li GL, Taschenberger H, Spirou G, Levinson SR, von Gersdorff H (2005) Presynaptic Na+ channels: locus, development, and recovery from inactivation at a high-fidelity synapse. J Neurosci 25:3724–3738.

Lorteije JA, Rusu SI, Kushmerick C, Borst JG (2009) Reliability and precision of the mouse calyx of Held synapse. J Neurosci 29:13770–13784.

Marom S, Levitan IB (1994) State-dependent inactivation of the Kv3 potassium channel. Biophys J 67:579–589.

Martinello K, Giacalone E, Migliore M, Brown DA, Shah MM (2019) The subthreshold-active KV7 current regulates neurotransmission by limiting spike-induced Ca(2+) influx in hippocampal mossy fiber synaptic terminals. Commun Biol 2:145.

Mc Laughlin M, van der Heijden M, Joris PX (2008) How secure is in vivo synaptic transmission at the calyx of Held? J Neurosci 28:10206–10219.

Peretz A, Pell L, Gofman Y, Haitin Y, Shamgar L, Patrich E, Kornilov P, Gourgy-Hacohen O, Ben-Tal N, Attali B (2010) Targeting the voltage sensor of Kv7.2 voltage-gated K+ channels with a new gating-modifier. Proc Natl Acad Sci U S A 107:15637–15642.

Peters HC, Hu H, Pongs O, Storm JF, Isbrandt D (2005) Conditional transgenic suppression of M channels in mouse brain reveals functions in neuronal excitability, resonance and behavior. Nat Neurosci 8:51–60.

Qi Y, Wang J, Bomben VC, Li DP, Chen SR, Sun H, Xi Y, Reed JG, Cheng J, Pan HL, Noebels JL, Yeh ET (2014) Hyper-SUMOylation of the Kv7 potassium channel diminishes the M-current leading to seizures and sudden death. Neuron 83:1159–1171.

Raman IM, Bean BP (2001) Inactivation and recovery of sodium currents in cerebellar Purkinje neurons: evidence for two mechanisms. Biophys J 80:729–737.

Sabatini BL, Regehr WG (1997) Control of neurotransmitter release by presynaptic waveform at the granule cell to Purkinje cell synapse. J Neurosci 17:3425–3435.

Sabatini BL, Regehr WG (1999) Timing of synaptic transmission. Annu Rev Physiol 61:521–542.

Schwarz JR, Glassmeier G, Cooper EC, Kao TC, Nodera H, Tabuena D, Kaji R, Bostock H (2006) KCNQ channels mediate IKs, a slow K+ current regulating excitability in the rat node of Ranvier. J Physiol 573:17–34.

Siegelbaum SA, Camardo JS, Kandel ER (1982) Serotonin and cyclic AMP close single K+ channels in Aplysia sensory neurones. Nature 299:413–417.

Sierksma MC, Borst JGG (2017) Resistance to action potential depression of a rat axon terminal in vivo. Proc Natl Acad Sci U S A 114:4249–4254.

Smith PH, Rhode WS (1987) Characterization of HRP-labeled globular bushy cells in the cat anteroventral cochlear nucleus. J Comp Neurol 266:360–375.

Soh H, Pant R, LoTurco JJ, Tzingounis AV (2014) Conditional deletions of epilepsy-associated KCNQ2 and KCNQ3 channels from cerebral cortex cause differential effects on neuronal excitability. J Neurosci 34:5311–5321.

Song P, Yang Y, Barnes-Davies M, Bhattacharjee A, Hamann M, Forsythe ID, Oliver DL, Kaczmarek LK (2005) Acoustic environment determines phosphorylation state of the Kv3.1 potassium channel in auditory neurons. Nat Neurosci 8:1335–1342.

Taschenberger H, von Gersdorff H (2000) Fine-tuning an auditory synapse for speed and fidelity: developmental changes in presynaptic waveform, EPSC kinetics, and synaptic plasticity. J Neurosci 20:9162–9173.

Wladyka CL, Kunze DL (2006) KCNQ/M-currents contribute to the resting membrane potential in rat visceral sensory neurons. J Physiol 575:175–189.

Zhang Y, Huang H (2017) SK Channels Regulate Resting Properties and Signaling Reliability of a Developing Fast-Spiking Neuron. J Neurosci 37:10738–10747.

